# Dopamine 2 receptor ablation from cholinergic neurons attenuates L-DOPA induced dyskinesias

**DOI:** 10.1101/2024.05.10.593604

**Authors:** Santiago Uribe-Cano, Lauren Malave, Andreas H. Kottmann

## Abstract

Striatal cholinergic interneurons (CIN) have been implicated in both, the facilitation as well as the attenuation of L-DOPA-induced dyskinesias (LID). These findings indicate that CIN impinge on formation and expression of LID in a dopamine state dependent manner since LID formation requires prominent oscillations of dopamine at timescales of hours over several years. However, how CIN sense and interpret striatal dopamine levels is not completely understood. CIN express both inhibitory, high affinity, Gαi coupled D2 (D2R)- and facilitatory, medium affinity, Gαs coupled D5 (D5R)-dopamine receptors. While the systemic ablation of D5R exacerbates LID, the contribution of D2R expression in CIN to LID has not been studied. Here, we produced mice with conditional ablation of D2R from choline acetyltransferase–expressing cells (D2_ChAT_KO) subjected to unilateral 6-hydroxydopamine lesions and chronic L-DOPA dosing. Behavioral assessments revealed that D2_ChAT_KO mice exhibited attenuated LID across escalating L-DOPA doses. Postmortem analyses showed reduced expression of the LID-associated marker p-ERK in CIN in the dorsolateral striatum. Further, quantification of the CIN activity marker p-rpS6^240/244^ of mice in the L-DOPA ON and OFF state revealed that L-DOPA resulted in an increase of cholinergic activity driven by a subset of mainly dorso-laterally located CIN. D2R ablation from CIN prevented the L-DOPA associated increase in cholinergic activity. Together, these findings indicate that D2R signaling in CIN promotes LID formation, and they highlight CIN D2R as a potential molecular target for mitigating dyskinesias while preserving the therapeutic efficacy of L-DOPA. We discuss our results in the context of recently refined models how CIN contribute to aberrant plasticity in the basal ganglia of mouse models of Parkinson’s Disease.

## 1 Introduction

Dopamine (DA) substitution therapy with the DA precursor L-3,4-dihydroxyphenylalanine (L-DOPA) remains the most efficacious treatment of the motor symptoms of Parkinson’s disease (PD) (Cotzias, Papavasiliou, and Gellene 1969; Albin, Young, and Penney 1989). However, most patients with advanced PD develop debilitating involuntary movements known as L-DOPA-induced dyskinesias (LID), which are caused by the slow oscillations of drug associated DA levels (de la Fuente-Fernandez et al. 2004; Bastide et al. 2015). LID formation and expression requires a hypodopaminergic state and consequent hypersensitivity to DA brought about by the underlying degeneration of DA neurons (DAN) in PD and can be modeled in rodents and non-human primates (Cenci and Lundblad 2007; Cenci, Jorntell, and Petersson 2018). These motor complications are largely attributed to aberrant synaptic plasticity of spiny projection neurons (SPN) in the striatum, the largest information input structure of the basal ganglia and a hub for motor learning (Cheung et al. 2023; Zhai et al. 2017; Redgrave et al. 2010). Under normal conditions, elevations of DA above tonic levels caused by its phasic release from mesencephalic projections to the striatum results in the reinforcement of future actions that were associated with positive outcomes in the past, while phasic reductions in DA below tonic levels results in the inhibition of actions in the future that were associated with negative outcomes in the past (Redgrave et al. 2010; Berke 2018). Consistent, DAN degeneration and resulting hypodopaminergia causes an impoverishment of movement over time through skewed motor learning that favors bradykinesia (Cheung et al. 2023). In contrast, the non-physiological, pulsatile surges of DA signaling produced by each L-DOPA dose uncoupled from a physiological evaluation of concurrent actions may reinforce context-inappropriate motor programs that ultimately produce dyskinesias (Bastide et al. 2015; Zhai et al. 2025; Cheung et al. 2023; Figge et al. 2024; Ryan et al. 2024; Girasole et al. 2018; Zhuang, Mazzoni, and Kang 2013).

In the healthy brain, DA action in the striatum is powerfully controlled by ACh, which is released by cholinergic interneurons. CIN are recognized for their tonic pacemaker-like activity in the 5–10 Hz range. In addition, CIN show complex phasic activity patterns in response to salient stimuli that consist of sustained pauses often flanked by bursts of spikes that lead to ACh fluctuations in the dorso lateral striatum (Aosaki et al. 1994; Apicella et al. 2011; Morris et al. 2004; Kimura, Rajkowski, and Evarts 1984; Ravel, Legallet, and Apicella 1999; Schulz and Reynolds 2013; Zhang and Cragg 2017; Krok et al. 2023; Duhne et al. 2024). These cholinergic pauses develop during reinforcement learning in a manner anticorrelated with striatal DA release (Morris et al. 2004; Duhne et al. 2024). The molecular mechanisms underlying the “conditioned” pause have received considerable attention, in part because the resulting reductions in ACh are thought to create a temporal window during which DA can modify corticostriatal synapses on SPN (Kimura, Rajkowski, and Evarts 1984; Aosaki, Graybiel, and Kimura 1994; Graybiel et al. 1994; Morris et al. 2004; Wang et al. 2006). Glutamatergic input strength, intrinsic electrophysiological properties, GABAergic inhibition, and DA signaling have all been implicated in shaping CIN pauses in the intact brain (reviewed in (Ratna and Francis 2025). Moreover, repeated DAN stimulation drives progressive shortening of the CIN pause in mice (Uribe-Cano and Kottmann 2025) suggesting that CIN themselves undergo DA mediated long term plasticity.

In DA depleted animals, repeated L-DOPA bolus treatment distorts normal CIN activity, both acutely and long term, leading to elevated tonic firing, enhanced GABAergic input, abolishment of the pause and disturbed learning (Choi et al. 2020; Aosaki et al. 1994; Aosaki, Graybiel, and Kimura 1994; Tubert et al. 2025; Cheung et al. 2023; Zhai et al. 2025). Whether and how these changes in CIN are drivers or associations of LID remain incompletely understood (Shen, Zhai, and Surmeier 2022; Bordia et al. 2016; Lim, Kang, and McGehee 2014). For example, boosting as well as blocking cholinergic signaling and cyto-toxic ablation of CIN can attenuate LID (Zhai et al. 2025; Nielsen and Ford 2024; Shen et al. 2016) (Ding et al. 2011; Won et al. 2014). Nonetheless, there is no doubt that L-DOPA impacts CIN activity acutely and long-term and that CIN activity profoundly affects LID.

CIN cell-autonomous sensitivity to DA, and therefore to L-DOPA, is largely mediated by dopamine D2R and D5R, suggesting that the pathological activity of CIN induced by L-Dopa may at least in part be caused by altered signaling through one or both of these receptors (Yan, Song, and Surmeier 1997; Jarvie et al. 1993; Tiberi and Caron 1994; Tiberi et al. 1996) supporting this possibility, prior studies indicated that the systemic ablation of the medium affinity and high constitutive activity dopamine D5R (Tiberi et al. 1991), a Gαs-coupled GPCR (Grandy et al. 1991), facilitates LID (Castello et al. 2020). Specifically, D5R knockout mice displayed exacerbated abnormal involuntary movements (AIMs), which worsened with repeated L-DOPA dosing. Although these studies relied on global ablation of D5R, the restricted expression of D5R in other striatal cell types lend support to a CIN-specific mechanism (Yan and Surmeier 1997). Consistent, a recent study found evidence that inverse agonism of D5 by clozapine can restore the cholinergic pause in L-DOPA treated parkinsonian mice by relieving of Kv1 channels from tonic D5R inhibition in CIN (Tubert et al. 2025). In contrast, the role of Gαi-coupled, high affinity dopamine D2R on CIN in LID remains less defined. Consistent with its Gαi coupling, D2R activation impinges on multiple intrinsic conductances in CIN and inhibits the autonomous spiking of CIN as well as ACh release from axon terminals (Chantranupong et al. 2023; Chuhma et al. 2014; Wieland et al. 2015; Straub et al. 2014; Ding et al. 2006). Further, cholinergic neuron specific loss and gain of function studies reveal that D2R signaling modulates the duration of the cholinergic pause in a graded manner (Kharkwal et al. 2016; Gallo et al. 2022; Martyniuk et al. 2022). This modulation of CIN by D2R is critical for the manifestation of dopamine dependent striatal plasticity (Wang et al. 2006).

Given the opposing regulation of CIN physiology by D2R and D5R, we hypothesized that D2R signaling in CIN might be a molecular target by which L-DOPA facilitates LID. Consistent with this idea, ablation of D2R from ChAT^+^ cells (D2_ChAT_KO) attenuated LID and prevented expression of the LID-associated marker p-ERK in CIN in the 6-OHDA model of LID. D2_ChAT_KO also blocked L-DOPA-associated increases in CIN activity markers—an outcome counterintuitive for the loss of an inhibitory receptor but aligned with prior evidence of heightened CIN activity during LID. Together, our findings suggest that removal of D2R from CIN might result in elevations of ACh that match L-DOPA dependent increases of DA signaling on SPNs in the DAN lesioned brain.

## 2 Materials and Methods

### 2.1 Anesthesia

Anesthesia was induced using a gas mixture of oxygen (1.5 l/min) and 4% isoflurane (Covetrus).

### 2.2 Euthanasia

Euthanasia was achieved by terminal transcardial perfusion. Mice were deeply anesthetized with intraperitoneal pentobarbital (15 mg/kg) and perfused transcardially with 0.9% saline followed by 4% paraformaldehyde (PFA) in 0.1 M phosphate-buffered saline (PBS).

### 2.3 Animals

Cholinergic-specific allele recombination was driven by Cre expression under the control of the choline acetyltransferase promoter (ChATCre^+/-^) (Rossi et al. 2011). Conditional ablation of D2R was achieved using a floxed D2R allele (Bello et al. 2011). Experimental animals were generated from a D2^L/+^:ChATCre^+/-^ × D2^L/+^ cross, following an initial ChATCre^+/-^ × D2^L/L^ pairing. Both sexes were included and balanced across conditions. Homozygous D2^L/L^:ChATCre^+/-^ animals (D2_ChAT_KO) lacked D2R in ChAT^+^ cells, among which CIN are included. Controls were ChATCre^+/-^ or heterozygous D2^L/+^:ChATCre^+/-^mice.

### 2.4 6-OHDA Lesion and L-DOPA Treatment

To induce a Parkinsonian state, mice received unilateral intrastriatal 6-OHDA in the dorsolateral (DL) striatum under anesthesia induced using a gas mixture of oxygen (1.5 l/min) and 4% isoflurane (Covetrus). Specifically, two unilateral injections (2 × 2µl each) of 6-OHDA were administered into the left striatum at coordinates anteroposterior (AP): +1.0mm, mediolateral (ML): +2.1mm, dorsoventral (DV): −2.9mm; and AP: +0.3mm, ML: +2.4mm, DV: −2.9mm. Injections were performed using a Hamilton syringe attached to a micro-syringe pump at a rate of 0.4μl/min. The syringe was left in place for 3 minutes following infusion and then retracted slowly to minimize backflow. A minimum of 18 days was allowed post-surgery to ensure sufficient degeneration of DAN before onset of behavioral testing.

After the recovery period, Parkinsonian phenotype was confirmed via open-field turn bias and cylinder paw-use tests. Open field testing was performed in a 43 × 43cm open-field arena (ENV 515S; Med Associates), and cylinder tests were performed in a large 2L glass beaker.

L-DOPA (5–20Mg/Kg) was administered daily in the home cage, except on days when behavioral testing occurred. Therapeutic response was assessed after the first dose using the open-field and cylinder tests. L-DOPA dosing escalated as follows: 5Mg/Kg (Days 1–13), 10Mg/Kg (Days 14–16), 15Mg/Kg (Days 17–18), and 20Mg/Kg (Days 19–31). After 31 days, half the animals received 9g/l saline for one week (washout), while the other half continued L-DOPA treatment (20Mg/Kg). Animals were sacrificed 30 minutes after the final dose for immunohistochemistry. All behavioral testing was performed by experimenters blinded to genotype. Mice with less than 20% reduction of the dopaminergic marker Tyrosine Hydroxylase (TH) in the lesioned hemisphere were excluded (2 controls, 1 mutant).

### 2.5 Dyskinesia Quantification

Abnormal involuntary movements (AIMs) were regularly quantified throughout the dosing regimen. On assessment days, animals received the specified L-DOPA dose offset from each other by one-minute intervals and were placed individually in empty housing cages without bedding. At 20-minute intervals, over a total duration of two hours, each animal was serially observed for a one-minute period. During each observation, the total number of contralateral rotations and the severity of AIMs were recorded in accordance with validated methods (Sebastianutto et al. 2016a, 2016b). AIM severity was scored for each of the defined subtypes (axial, limb, orofacial) at each one-minute interval on a scale of 0–4: 0 = no expression of the subtype; 1 = expression for less than 50% of the observation period; 2 = expression for more than 50% of the observation period; 3 = continuous but interruptible expression throughout the observation period; 4 = continuous and uninterruptible expression throughout the observation period. AIM scores across the two-hour session were summed for each mouse to yield a total AIM score. Data were collected from 3 independent cohorts: 1 utilizing heterozcygous D2^L/+^:ChATCre^+/-^ controls and 2 utilizing ChATCre^+/-^controls.

### 2.6 Immunohistochemistry

Mice were deeply anesthetized with intraperitoneal pentobarbital (15 mg/kg) and perfused transcardially with 0.9% saline followed by 4% paraformaldehyde (PFA) in 0.1 M phosphate-buffered saline (PBS). Brains were post-fixed overnight at 4°C in 4% PFA, then cryoprotected in 30% sucrose in PBS until they sank (∼48 hours). Cryoprotected brains were embedded in optimal cutting temperature (OCT) compound, frozen on dry ice, and stored at −80°C.

Brains were cryo-sectioned coronally at 30 µm using a cryostat at −20°C and stored as free-floating sections in PBS with 0.1% sodium azide until staining.

Anti-tyrosine hydroxylase (1:500; RRID: AB_657012) and anti-ChAT (1:100; RRID: AB_144P) were purchased from Millipore. Anti-phospho-Erk1/2 (1:400; RRID: AB_9101), and anti-phospho-rpS6 (ser240/244; RRID: AB_5364) were from Cell Signaling Technology. Anti-NeuN (1:200; RRID: AB_MAB377) from Chemicon International. Various Alexa Fluor antibodies were used at 1:250 dilutions for immunohistochemistry (Jackson Immuno Research).

Images of fluorescent immunoreactivity used for quantification were acquired using a ZEISS LSM 880 confocal microscope. Most images were taken at 20× magnification (10× for assessing 6-OHDA-induced DAN axon loss). Confocal fluorescence was quantified via ImageJ. For DAN loss, TH Mean Gray Value (MGV) was measured in the dorsal striatum and normalized to background. Percent TH was calculated relative to the unlesioned hemisphere. Lesion extent was mapped across multiple striatal/midbrain sections (Supplementary Figure 1). For p-ERK and p-rpS6^240/244^ analysis, 700um^2^ fields centered at DL (ML: +/-2.2mm; DV: −3.0mm) or dorsomedial (DM) (ML: +/-1.2mm; DV: −2.5mm) positions and spanning AP: 0.3-1mm were used.

For p-ERK, MGV was measured in CIN (ChAT^+^ cells) of both hemispheres. CIN were considered p-ERK^+^ if their MGV exceeded the unlesioned hemisphere mean +2 SD. Percentage of p-ERK^+^ CIN was reported per lesioned striatum. The same procedure was used for p-ERK^+^/NeuN^+^ cells (excluding ChAT^+^ cells).

For p-rpS6^240/244^, MGV was calculated for ChAT^+^ regions of interest (ROIs) and normalized to NeuN MGV within the same ROI.

## 3 Results

### 3.1 Validation of D2_ChAT_KO 6-OHDA Model

To selectively ablate the D2R from CIN, we generated mice homozygous for a floxed D2 allele (D2^L/L^) and expressing Cre recombinase under the choline acetyltransferase (ChAT) promoter (D2^L/L^:ChATCre^+/-^). Littermate controls heterozygous for the floxed allele and carrying the ChATCre driver (D2^L/+^:ChATCre^+/-^) were used to control for both Cre expression and the introduction of the floxed allele. Successful recombination of the D2R locus was confirmed in striatal, but not tail, tissue (**Figure 1A**).

**Figure 1.**
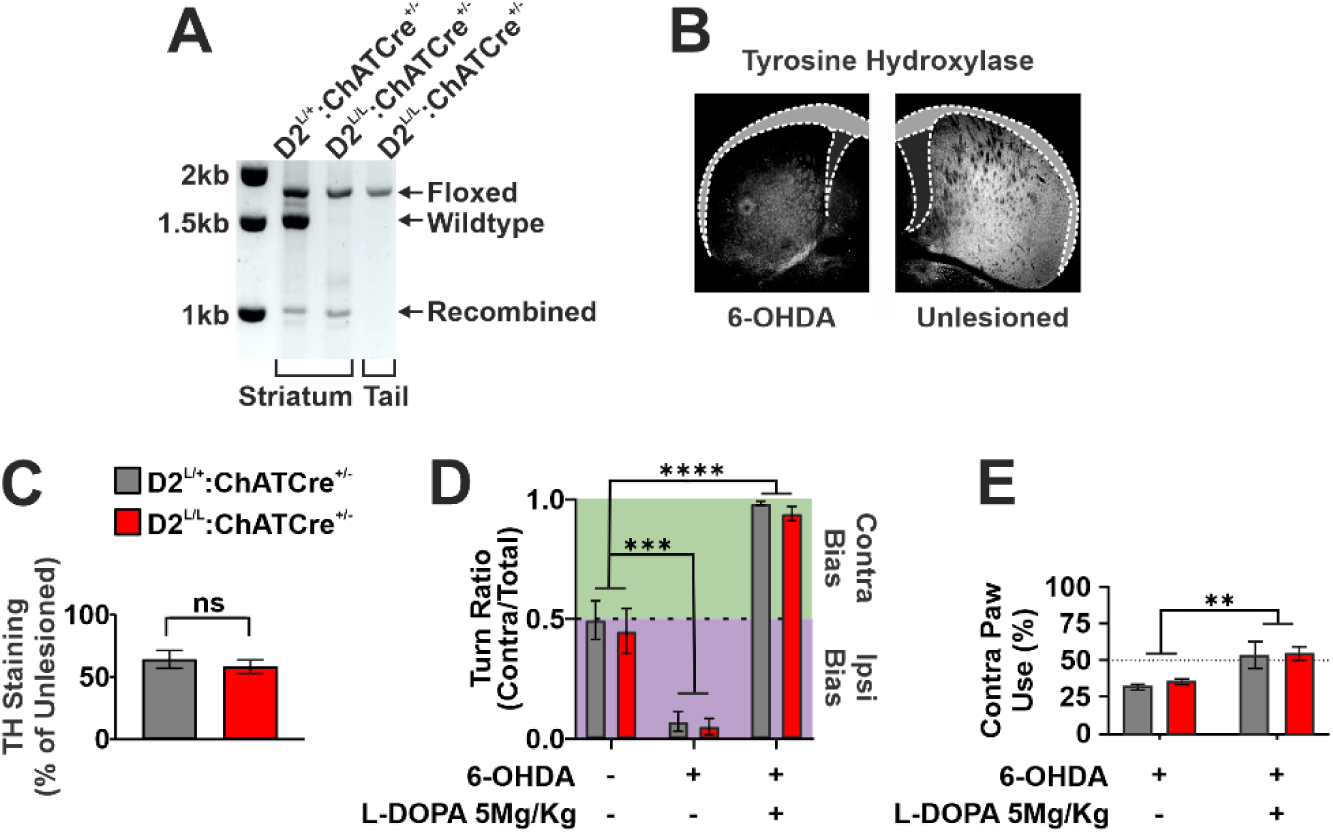
**(A)** PCR validation of ChATCre-dependent recombination of the dopamine receptor 2 (D2) allele. Primers amplify regions spanning exon 2 and the flanking LoxP sites. Absence of a recombined band in tail samples confirms recombination is restricted to ChATCre-expressing cells, including cholinergic interneurons (CIN) of the striatum. **(B)** Representative image showing tyrosine hydroxylase (TH) depletion in the dorsal striatum following 6-hydroxydopamine (6-OHDA) injection. **(C)** Quantification of TH depletion in the lesioned dorsal striatum, expressed as the lesioned/unlesioned hemisphere ratio, shows no difference between control and D2^L/L^:ChATCre^+/-^ animals (n = 5–7 per genotype; unpaired two-tailed Student’s t test, p > 0.05). **(D)** Turn bias in the open field, measured at baseline, after unilateral 6-OHDA lesion, and after subsequent L-DOPA treatment (5Mg/Kg). Turn bias was quantified as the ratio of contralateral turns over total turns. Both genotypes showed comparable reversal of 6-OHDA-induced turn bias following L-DOPA treatment (n = 5–7 per genotype; mixed-effects model: Timepoint effect, F(1.220, 17.69) = 109.7, p < 0.0001; Genotype effect, F(1, 29) = 0.55, p > 0.05; Timepoint × Genotype interaction, F(2, 29) = 0.03, p > 0.05; post hoc Tukey’s multiple comparisons test: ***p < 0.001, ****p < 0.0001). **(E)** Cylinder test measuring the percentage of paw rears performed with the forelimb contralateral to the lesion, before and after L-DOPA treatment. Both genotypes exhibited similar L-DOPA-dependent increases in contralateral paw use (n = 5–7 per genotype; two-way repeated-measures ANOVA: Timepoint effect, F(1, 10) = 20.67, p < 0.01; Genotype effect, F(1, 10) = 0.21, p > 0.05; Timepoint × Genotype interaction, F(1, 10) = 0.08, p > 0.05).

Following intrastriatal 6-OHDA injections targeting the dorsolateral (DL) striatum, both D2^L/L^:ChATCre^+/-^ and control animals exhibited similar levels of striatal dopamine neuron (DAN) degeneration, indicating that D2R deletion from CIN does not alter 6-OHDA-induced cytotoxicity (**Figure 1B, C).**

To validate the Parkinsonian phenotype, we employed two standard assays of unilateral motor dysfunction. In the open field, 6-OHDA lesions induced a robust ipsilateral turning bias (**Figure 1D**). Similarly, the cylinder test revealed an ipsilateral forelimb-use preference, confirming equivalent hypodopaminergic motor deficits across genotypes (**Figure 1E**).

Subsequent administration of 5Mg/Kg L-DOPA reversed these motor asymmetries in both assays: turning bias in the open field shifted contralaterally, and forelimb use in the cylinder test similarly favored the contralateral paw (**Figure 1D, E**). These effects were equivalent in control and D2^L/L^:ChATCre^+/-^animals, indicating that deletion of D2 from CIN does not impair the prokinetic efficacy of L-DOPA.

### 3.2 Loss of D2 on ChAT^+^ Cells Attenuates L-DOPA Induced Dyskinesias

Treatment with L-DOPA over the course of the experiment consisted of a chronic dosing schedule that escalated over the course of 31 days, after which, half the animals from each genotype underwent a 1-week washout period during which they received saline while the other half continued L-DOPA dosing at the highest dose of 20Mg/Kg. The rationale for this design was twofold. First, p-ERK, a validated histological marker of LID severity, implicates CIN as critical contributors in later stages of LID (Ding et al. 2011). We therefore aimed to capture these progressive mechanisms with an extended dosing regimen. Second, we sought to assess how CIN, with or without D2R expression, responded to L-DOPA once LID was fully established. By isolating the contribution of D2R signaling to the acute induction of LID at this stage, we hoped to gain insight into the mechanisms that sustain LID once developed, independent of the early-stage plasticity likely prominent during initial L-DOPA treatment.

Control animals displayed persistent abnormal involuntary movements (AIMs) that worsened as L-DOPA dose increased on days 14, 17, and 19 (**Figure 2A**). In contrast, D2_ChAT_KO mice exhibited consistently attenuated AIMs, with the largest differences from controls observed at higher doses. Reductions across axial, limb, and orofacial AIM subtypes accounted for the lower total AIM scores in D2_ChAT_KO animals (**Figure 1B**).

**Figure 2.**
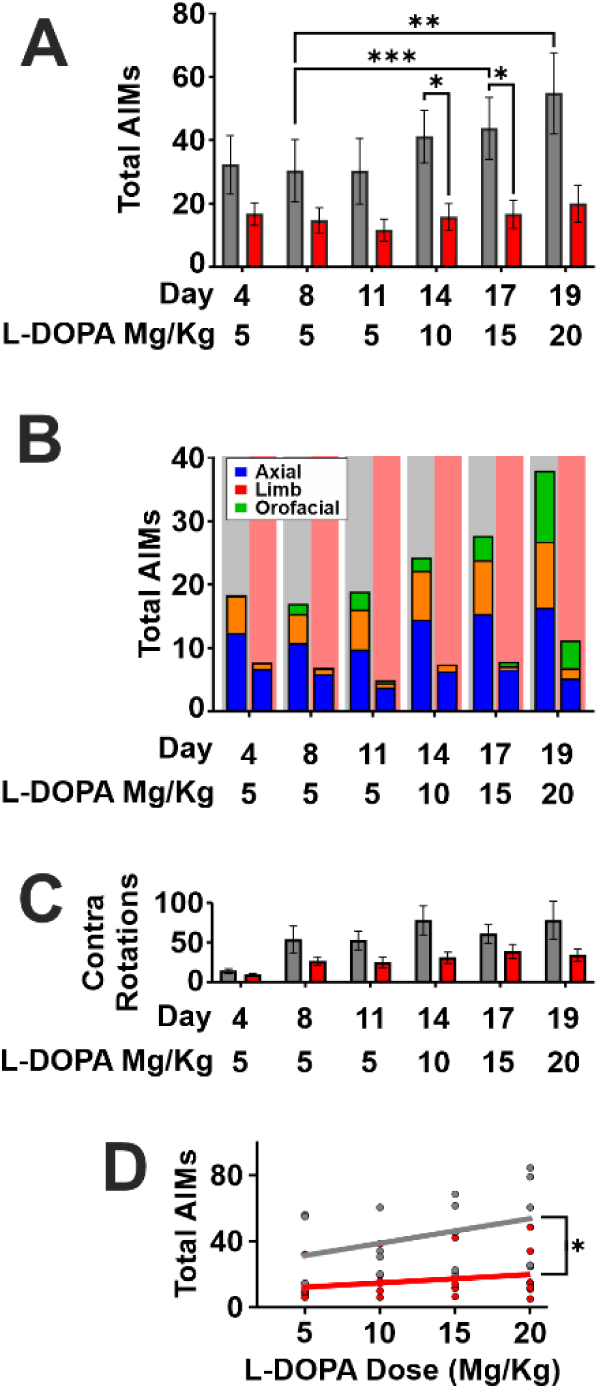
**(A)** Total abnormal involuntary movement (AIM) scores over the first 19 days of L-DOPA treatment (n = 5–7 per genotype; two way repeated-measures ANOVA: Day/Dose effect, F(2.462, 24.62) = 11.29, p < 0.001; Genotype effect, F(1, 10) = 6.03, p < 0.05; Genotype × Day/Dose interaction, F(5, 50) = 4.50, p < 0.01; post hoc Tukey’s multiple comparisons test: *p < 0.05, **p < 0.01, ***p < 0.001). **(B)** Breakdown of AIM subtypes across L-DOPA treatment days. Stacked bars represent the average group values for each AIM subtype; column background indicates genotype. **(C)** Total contralateral rotations during AIM scoring sessions across the first 19 days of L-DOPA treatment (n = 5–7 per genotype; two-way repeated-measures ANOVA: Day/Dose effect, F(2.311, 23.11) = 15.92, p < 0.0001; Genotype effect F(1, 10) = 4.75, p > 0.05; Genotype × Day/Dose interaction, F(5, 50) = 3.53, p < 0.01). **(D)** AIM severity plotted against L-DOPA dose for each genotype. Slope differences assessed by linear regression followed by ANCOVA (n = 5–7 per genotype; F(1, 4) = 15.97, p < 0.05).

When comparing total contralateral rotations between genotypes on AIM scoring days, we observed an interaction effect. While there was no difference in rotation bias in the open field on day 4 of L-DOPA treatment, control animals developed a progressive increase in rotational bias over time (**Figure 1C**).

LID severity is known to be associated with the strength of L-DOPA dose administered. To see if this correlation was impacted by D2R ablation from cholinergic neurons, we performed linear regression of total AIM severity across the four L-DOPA doses tested for both controls and D2_ChAT_KO animals. The resulting slopes differed significantly between genotypes, indicating that the positive correlation between L-DOPA dose and AIM severity observed in controls was attenuated among D2^L/L^:ChATCre^+/-^ animals. This analysis further supported the absence of LID escalation in D2^L/L^:ChATCre^+/-^ mice compared to controls (**Figure 1I**).

To address potential gene dosage effects in heterozygous controls, we replicated these behavioral experiments using pooled data from independent cohorts with alternative ChATCre^+/-^controls carrying two wild-type D2 alleles (**Supplementary Figure 1**). Consistent with the first cohort, control animals again exhibited persistent AIMs that scaled with L-DOPA dose, whereas D2^L/L^:ChATCre^+/-^ mice displayed attenuated dyskinesias across doses (**Supplementary Figure 1D, G).** To address the possibility of reduced DAN lesioning by 6-OHDA in D2^L/L^:ChATCre^+/-^ mice we performed detailed mapping of tyrosine hydroxylase (TH) loss in the striatum and midbrain of both genotypes(**Supplementary Figure 2**). This analysis revealed substantial reduction in DAN neurite density in the striatum and corresponding DAN cell body density in the 6-OHDA-treated hemispheres of both mutants and controls. These results rule out that the observed differences in LID severity are caused by differential lesion extent or severity between groups.

### 3.3 Reduced CIN Expression of p-ERK Reflects Attenuated AIMs among D2_ChAT_KO Animals

Phosphorylation of extracellular signal-regulated kinase1/2 (p-ERK) in the hypodopaminergic striatum reflects LID severity and progression (Malave et al. 2021; Won et al. 2014; Ding et al. 2011). Rodent models of LID have shown that p-ERK expression occurs in both MSN and CIN and is correlated with severity of dyskinesias. Notably, early-stage L-DOPA treatment primarily increases p-ERK in MSN, while chronic L-DOPA exposure shifts this expression to CIN (Won et al. 2014). Moreover, pharmacological inhibition of p-ERK has been shown to attenuate LIDs, suggesting that p-ERK activation may not only reflect, but also contribute to LID pathophysiology (Ding et al. 2011). We therefore assessed whether the differences in LID severity between heterozygous controls and D2^L/L^:ChATCre^+/-^ animals were reflected in p-ERK expression at the end of our chronic L-DOPA dosing regimen.

Consistent with the behavioral findings, D2^L/L^:ChATCre^+/-^ animals exhibited a significant reduction in p-ERK^+^ CIN (**Figure 3A, B**). While the interaction effect between genotype and striatal region only approached significance (p = 0.069), exploratory post hoc comparisons nevertheless revealed a significant genotype difference specifically in the DL striatum, an effect consistent with the localized impact of our DL 6-OHDA lesions and the DL striatum’s predominant role in LID formation and expression (**Girasole et al. 2018)**.

**Figure 3.**
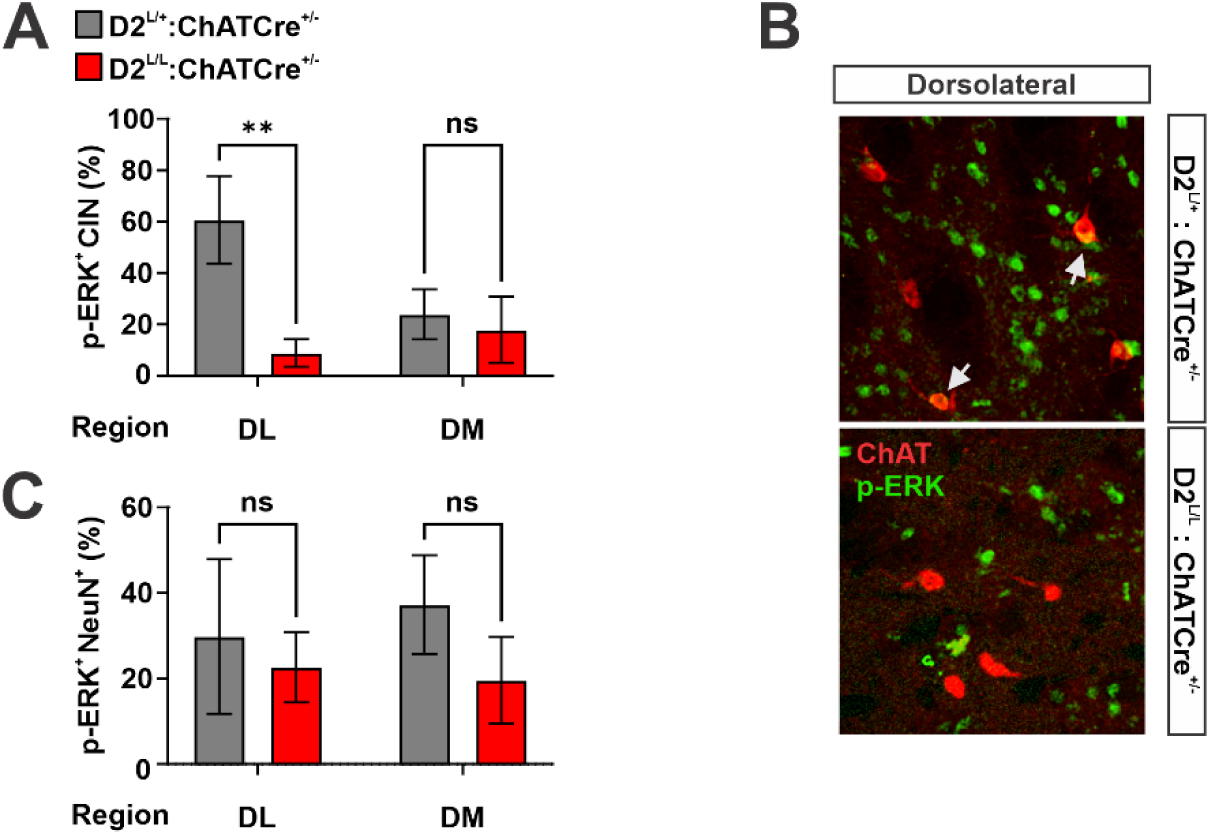
**(A)** Quantification of phosphorylated extracellular signal-regulated kinase 1/2 positive (p-ERK^+^) cholinergic interneurons (CIN), reported as the percentage of total CIN in the dorsolateral (DL) or dorsomedial (DM) striatum (n = 2–3 striatal sections per animal, across 2–3 animals per genotype; two-way repeated-measures ANOVA: Genotype effect, F(1, 12) = 5.96, p < 0.05; Region effect, F(1, 12) = 1.45, p > 0.05; Genotype × Region interaction, F(1, 12) = 3.96, p = 0.069; post hoc Šídák’s multiple comparisons test: **p < 0.01). **(B)** Representative images of p-ERK and choline acetyltransferase (ChAT) immunoreactivity in the DL striatum. **(C)** Quantification of p-ERK^+^/ChAT^-^ neurons, reported as the percentage of total ChAT^-^ neurons in the DL or DM striatum (n = 2–3 striatal sections per animal, across 2–3 animals per genotype; two-way repeated-measures ANOVA: Genotype effect, F(1, 12) = 0.05, p > 0.05; Region effect, F(1, 12) = 0.85, p > 0.05; Genotype × Region interaction, F(1, 12) = 0.27, p > 0.05).

When we examined p-ERK expression in non-cholinergic NeuN^+^ neurons of the striatum, which are mostly SPN, we found no genotype differences, consistent with the chronic nature of the L-DOPA dosing paradigm and the cholinergic neuron specificity of D2R ablation (**Figure 3C**).

### 3.4 D2_ChAT_KO prevents L-DOPA-Induced Increases in CIN-specific p-rpS6^240/244^

To assess D2R’s impact on CIN activity during LID we quantified levels of p-rpS6^240/244^, a CIN activity marker (**Figure 4A**) (Matamales, Gotz, and Bertran-Gonzalez 2016; Bertran-Gonzalez et al. 2012). Focusing on the established LID state, we compared DL CIN in control and D2_ChAT_KO mice following continued L-DOPA treatment (L-DOPA ON) or after a 1-week washout period (L-DOPA OFF).

**Figure 4.**
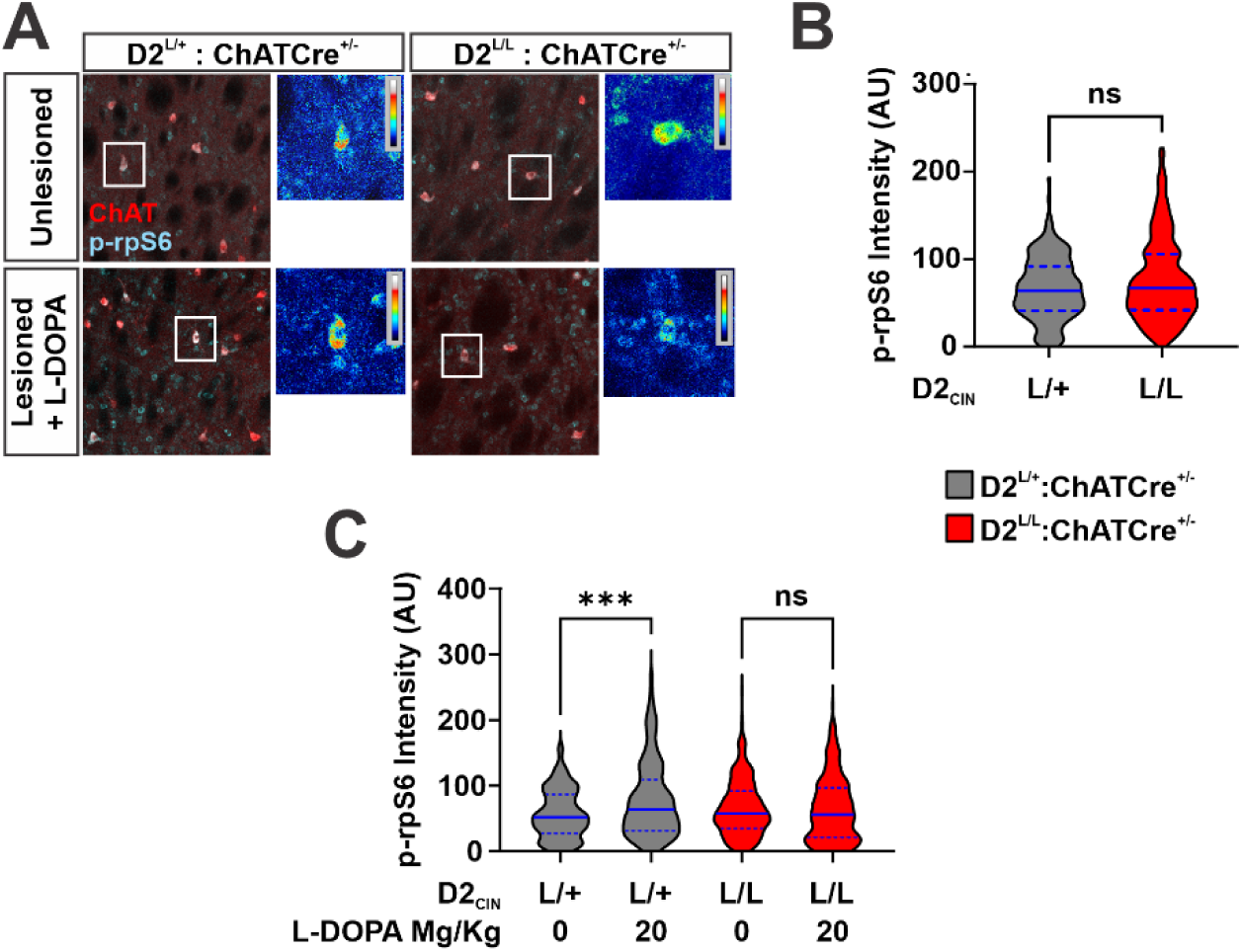
**(A)** Representative images from each genotype showing phosphorylated ribosomal protein S6 (p-rpS6^240/244^) and choline acetyltransferase (ChAT) immunoreactivity in the dorsolateral (DL) striatum under two conditions: unlesioned hemispheres (no 6-OHDA, L-DOPA washout) and lesioned hemispheres following L-DOPA treatment. White rectangles indicate inset regions displaying heat maps of p-rpS6^240/244^ intensity within individual CIN. **(B)** Quantification of normalized CIN p-rpS6^240/244^ intensity in the DL striatum of unlesioned hemispheres (no 6-OHDA, L DOPA washout; n = 150+ CIN sampled from 4 striatal sections per animal, across 2–3 animals per genotype; unpaired two tailed Student’s t test, p > 0.05). **(C)** Quantification of normalized CIN p-rpS6^240/244^ intensity in the DL striatum of 6-OHDA-lesioned hemispheres with and without L-DOPA washout (n = 150+ CIN sampled from 4 striatal sections per animal, across 2–3 animals per genotype; two way ANOVA: Genotype effect, F(1, 957) = 0.26, p > 0.05; L-DOPA effect, F(1, 957) = 8.29, p < 0.01; Genotype × L-DOPA interaction, F(1, 957) = 11.90, p < 0.001; post hoc Fisher’s LSD: ***p < 0.001).

In the unlesioned hemispheres of L-DOPA OFF animals, CIN p-rpS6^240/244^ levels were similar between genotypes, indicating no baseline effect of D2_ChAT_KO (**Figure 4B**). Likewise, CIN of the lesioned hemispheres in L-DOPA OFF animals showed no genotype difference (**Figure 4C; columns 1 vs 3**). However, in lesioned hemispheres of L-DOPA ON animals, CIN p-rpS6^240/244^ levels increased in controls but not D2_ChAT_KO mice (**FIG. 2F; columns 2 vs 4**). Given the well established Gai mediated inhibitory influence of D2R signaling on CIN. this finding suggests that L-DOPA increases inhibition of CIN in the absence of D2R compared to controls in the hypodopaminergic DL striatum.

## 4 Discussion

Our findings suggest that ablation of Gαi coupled D2R from cholinergic neurons attenuates LID following unilateral 6-OHDA lesioning and repeated bolus L-DOPA treatment. The results presented here not only complement previous studies which suggested that the Gαs coupled D5R from CIN facilitates LID but also might provide new insights into how DA modulation of CIN occurs in the healthy brain.

Multiple studies have shown that D2R agonism prolongs CIN afterhyperpolarizations, commonly referred to as the “conditioned pause”. In physiological contexts, phasic elevations of DA extends pause duration during behaviorally relevant striatal processing(Gallo et al. 2022; Martyniuk et al. 2022). More specifically, current motor learning models posit that the anticorrelation of elevated DA and reduced ACh creates a permissive coincidence window during which DA driven plasticity can occur at cortico-striatal synapses on striatal output neurons (SPN) (Deffains and Bergman 2015; Wieland et al. 2015) (Fig. 5A). Consistent, out of context, unphysiological L-Dopa surges are known to cause inappropriate plasticity leading to a randomization of synaptic weights and the appearance of un-purposeful movements (Fig. 5B,C) (Figge et al. 2024; Zhai et al. 2025). However, how dopaminergic control of CIN is involved in bringing about aberrant plasticity remains controversial and incompletely understood.

**Figure 5:**
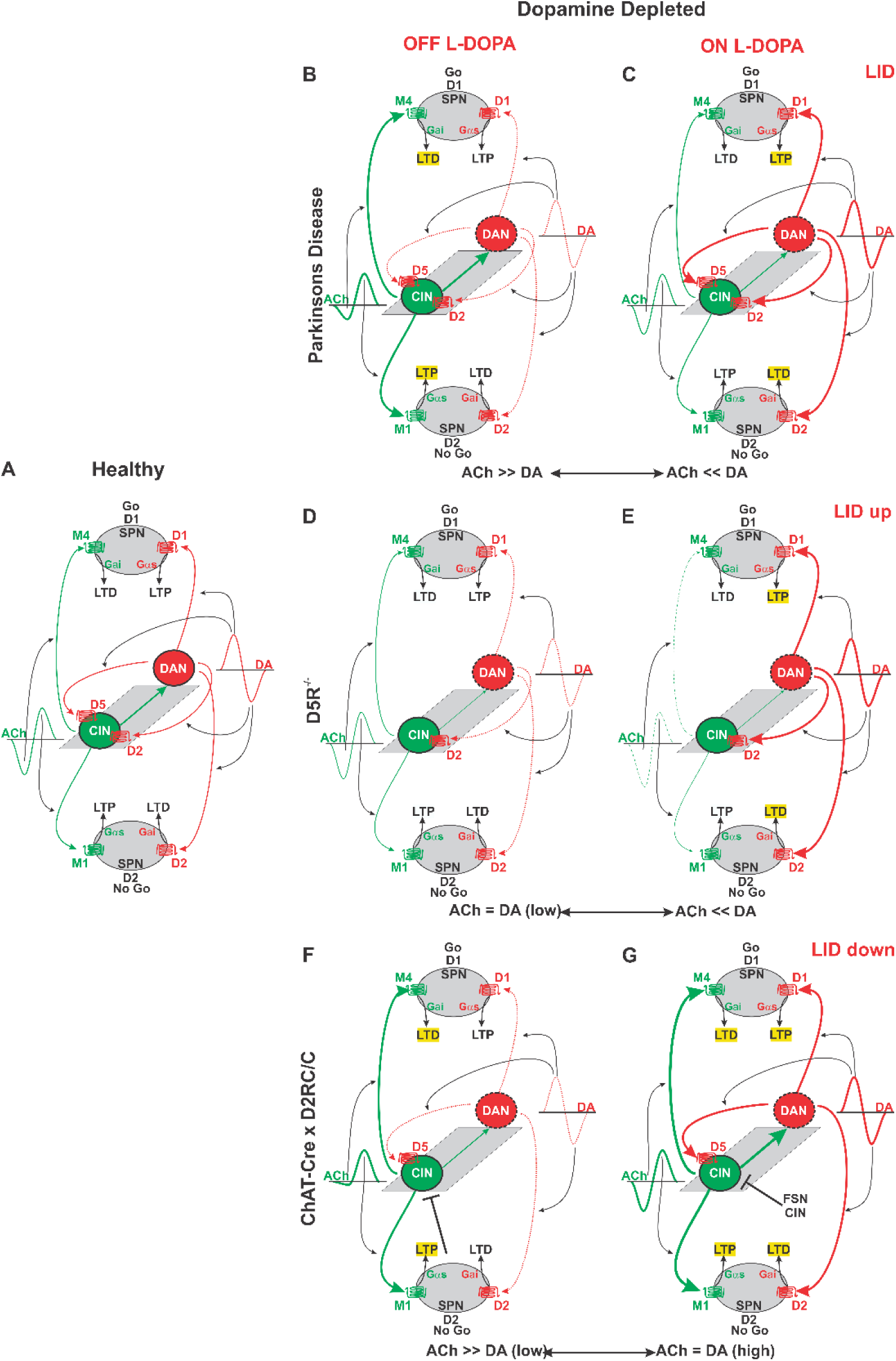
Model of reciprocal cholinergic (green) and dopaminergic (red) signaling. DAN degeneration indicated by dotted black line. DAN degeneration diminished dopaminergic signaling indicated by dotted red line.

### 4.2 Combined Effects of D2 and D5 Signaling on CIN in LID

Prior evidence from germline D5R ablation mice suggested that loss of D5R signaling enhances LID severity (Castello et al. 2020), whereas our findings presented here indicate that loss of D2R from CIN attenuates LID. These results support the idea that D2R and D5R exert opposing influences on CIN function in accordance with their coupling to either inhibitory Gαi or facilitatory Gαs, resp., during dyskinesia. However, the downstream effects on CIN physiology are less straightforward.

For instance, D5R ablation reduced p-ERK expression in CIN despite exacerbating LID, while D2R ablation also prevented p-ERK induction but with opposing effects on LID. Similarly, although D5R ablation decreased the CIN activity marker p-rpS6^240/244^ in line with its facilitatory role (Castello et al. 2020), D2R ablation likewise blocked its increase, suggesting that cellular changes brought about by the dyskinetic parkinsonian state—and its interaction with differences in receptor affinity or expression—may induce non-obvious and/or paradoxical receptor functions under certain conditions. How could these observations be integrated into the emerging model in which anti correlated ACh and DA fluctuations dynamically control reciprocally opposing activity of SPNs during motor learning and execution (**Figure. 5A**)?

SPNs, which constitute ∼90% of all striatal neurons, fall into two groups. “Direct” pathway SPNs (dSPNs) promote action selection and execution (Redgrave et al. 2010). dSPNs are positively modulated by DA via Gαs/olf coupled dopamine D1 receptors (D1R), which facilitate long term synaptic potentiation (LTP) of their glutamatergic input (Gerfen and Surmeier 2011; Zhai et al. 2019; Tritsch, Ding, and Sabatini 2012; Tritsch and Sabatini 2012; Surmeier, Carrillo-Reid, and Bargas 2011; Surmeier et al. 2007) (Figure 1A). Conversely, “indirect” pathway SPNs (iSPNs) suppress contextually inappropriate actions (Redgrave et al. 2010). iSPNs are negatively modulated by DA via Gαi coupled dopamine D2 receptors (D2R) attenuating glutamatergic transmission, and promoting long-term synaptic depression (LTD) (Gerfen and Surmeier 2011; Zhai et al. 2019) (Tritsch, Ding, and Sabatini 2012; Tritsch and Sabatini 2012; Surmeier, Carrillo-Reid, and Bargas 2011; Surmeier et al. 2007) (**Figure 5A**). Dopamine’s effects on SPNs are powerfully counteracted and balanced by acetylcholine (ACh) signaling. In dSPNs, which primarily express Gαi-coupled M4 muscarinic receptors (M4Rs), ACh signaling blunts the effects of D1R activation and promotes LTD induction (Longo et al. 2023; Shen et al. 2016; Laverne et al. 2022; Zhai et al. 2023). In contrast, in iSPNs, which only express Gαs coupled M1 muscarinic receptors (M1Rs), ACh signaling opposes the effects of D2R inhibition and instead enhances somatic excitability, dendritic integration, and LTP induction (Crittenden et al. 2021; Day et al. 2008; Shen et al. 2007; Oldenburg and Ding 2011). In the healthy brain, DA and ACh levels fluctuate in a behavioral context dependent manner, and, therefore, these reciprocal and converse effects of DA and ACh on direct and indirect SPNs provide high contrast means for the dynamic coordination of acute motor output and the modulation of long-term plasticity processes that refine future behavior through learning (**Figure. 5A**).

In the parkinsonian brain the degeneration of DAN results in severely reduced DA signaling (**Figure 5B**, dotted red lines) and a relative increase in ACh signaling (**Figure 5B**, thickened green lines). The relative increase in cholinergic tone is associated with increased excitability of CIN and a near loss of the cholinergic pause (Aosaki et al. 1994; Zhai et al. 2025). Reduced DA signaling significantly contributes to this increased CIN activity via D2R and D5R. Reduced D2R signaling results in relief of inhibition via GαI (Chantranupong et al. 2023; Chuhma et al. 2014; Straub et al. 2014; Wieland et al. 2014; Ding et al. 2010), while absence of ligand binding to D5R allows ligand independent, constitutive receptor activity and upregulation of CIN via Gαs (Tubert et al. 2025). The shift towards greater ACh tone favors LTD in D1SPNs, LTP in D2SPNs and akinesia, and results in a loss of skilled movement through disturbed motor learning (Cheung et al. 2023). In the PD treated striatum, L-DOPA increases dopaminergic tone relative to ACh in part by reinstating D2R mediated inhibition of CIN biasing signaling towards LTP in D1SPNs, LTD in D2SPNs, and movement facilitation (Choi et al. 2020; Aosaki et al. 1994; Aosaki, Graybiel, and Kimura 1994; Tubert et al. 2025; Cheung et al. 2023) (**Figure 5C**). Importantly, the alternation between the L-DOPA on and off states significantly contributes to the formation of LID (Zhai et al. 2025) providing a possible rationalization of how both, ablation of CIN (during the OFF state) and facilitating cholinergic signaling (during the ON state), could inhibit LID.

The absence of D5R from CIN dampens CIN activity in the DAN lesioned striatum (**Figure 5D**) as evidenced by reduced p-rpS6 and pERK levels in CIN in D5R knock out mice (Castello et al. 2020) and recordings of slices with DAN lesions and inhibition of the constitutive activity of D5R, which reinstates the pause by restoring Kv1 currents (Tubert et al. 2025). In the L-DOPA ON state the restored pause would be predicted to be further exaggerated by reinstated D2R mediated inhibition (**Figure 5E**). Alternations between the OFF and ON state, in conjunction with expanded CIN pauses in the ON stage would therefore bias striatal physiology towards exaggerated D1SPN LTP and D2SPN LTD providing a possible mechanism for the observed enhancement of LID formation and expression in D5R knockout mice compared to controls (Castello et al. 2020).

Within this framework, D2R ablation from CIN in the DAN lesioned striatum should have little additional effect during the L-DOPA OFF state compared to controls: constitutive D5R activity and absence of D2R inhibition should abrogate the cholinergic pause and drive up cholinergic basal activity (**Figure 5F**). As such, D2R ablation would be expected to favor D1SPN LTD (through M4R signaling), D2SPN LTP (via M1R signaling) and akinesia in the OFF state. In contrast to controls, however, in the ON state, DA binding to D5R will restore the pause and elevate cholinergic activity unimpeded by D2R mediated inhibition (**Figure 5G**). In this scenario, consistent with D2R’s critical involvement in maintaining the anticorrelation of DA and ACh fluctuations in the healthy striatum (Krok et al. 2023; Gallo et al. 2022; Martyniuk et al. 2022), D2R ablation from CIN would convert the anticorrelation of DA and ACh seen in controls to an increase of cholinergic tone concurrent with L-DOPA caused DA surges (**Figure 5G**). Together, these effects should curtail L-DOPA driven D1SPN LTP and D2SPN LTD through balanced and complementary ACh signaling onto SPNs and result in attenuated LID formation and expression (**Figure 5G).**

Lastly, notwithstanding reports of paradoxical CIN activation by D2R signaling (Helseth et al. 2021; Pisani et al. 2006), the elevated CIN activity in D2R ablation mice might lead to increased, inhibitory feedback onto CIN from fast spiking GABAergic interneurons or recurrent inhibition among CIN (**Figure 5G**, black inhibitory arrow) (English et al. 2011; Sullivan, Chen, and Morikawa 2008; Koos and Tepper 2002; Faust et al. 2016), which could underpin the unintuitive finding that D2R ablation blunts L-DOPA associated CIN activation measured by p-rpS6 (**Figure 4**).

### 4.3 Translational Strategies

Our data provides first evidence that ablating D2R from cholinergic neurons attenuates LID severity and reduces LID associated molecular markers in CIN. Systemic pharmacological inhibition of D2R would come with unacceptable side effects. However, there are at least two pathways towards spatially and cell type selective D2R inhibition. The development of transcriptional cis acting elements that limit gene expression to CIN could be used for devising a CIN selective, local gene therapy strategy (Hunker et al. 2025). Another opportunity might arise from CIN cellular heterogeneity and the combinatoric influences of additional modulators of CIN activity. D2R signaling effects are greatest in the DL striatum (Chuhma et al. 2017; Chuhma et al. 2014), where LID associated neuronal ensembles can be identified (Girasole et al. 2018). Consistent, we find that the greatest effect of D2R ablation on CIN prp-S6 levels occurs at the lateral edge of the lesion area in the DL striatum (**Figure 4**). We previously found that stimulating the GPCR smoothened on CIN in the DL striatum attenuates LID, counteracts DA mediated inhibition of CIN and shortens the cholinergic pause (Malave et al. 2021; Uribe-Cano and Kottmann 2025). D2R and Smo are expressed in the primary cilium of CIN (Marley and von Zastrow 2010; Miyoshi et al. 2014; Lucarelli et al. 2019), which recently became implicated in the majority of sporadic PD (Khan et al. 2021; Khan et al. 2024; Nair et al. 2025; Uribe-Cano and Kottmann 2024). The likely close spatial proximity of these receptors in the primary cilium of CIN might make feasible the development of drugs that combine Smo agonist and D2R antagonist properties resulting in increased cellular selectivity and efficacy.

Although these findings and our rationalization attempts will require further validation, they highlight D2R in CIN as a molecular target for strategies aimed at preserving the therapeutic benefits of L-DOPA while limiting its debilitating motor side effects.

## Acknowledgments

We thank A. Walls for helpful discussion and technical assistance with PCR assays. We also thank K. Liu and M. Holmes for assistance with tissue preparation, staining, and data collection. S.U.C. was supported by grant NIH T32GM136499 (PIs Ruth Stark and Mark Steinberg) and A.H.K. was supported by NIH NS095253 (PI Andreas H Kottmann), AG065682 (PI Andreas H Kottmann), American Parkinson’s Disease Association (APDA; PI Andreas H Kottmann) and NIH U54MD017979 (PI Maria Lima).

## Author Roles

1. Research project: A. Conception, B. Organization, C. Execution;

2. Statistical Analysis: A. Design, B. Execution, C. Review and Critique;

3. Manuscript Preparation: A. Writing of the first draft, B. Review and Critique;

Santiago Uribe-Cano: 1A, 1B, 1C, 2A, 2B, 3A, 3B

Lauren Malave: 1A, 2C, 3B

Andreas H Kottmann: 1A, 1B, 2C, 3A, 3B

All authors had final approval of the submitted vers**Disclosures:**

## Ethical Compliance Statement

- All mouse experiments were approved by the IACUC committee of City College of New York.

---

**Supplementary Figure 1.**
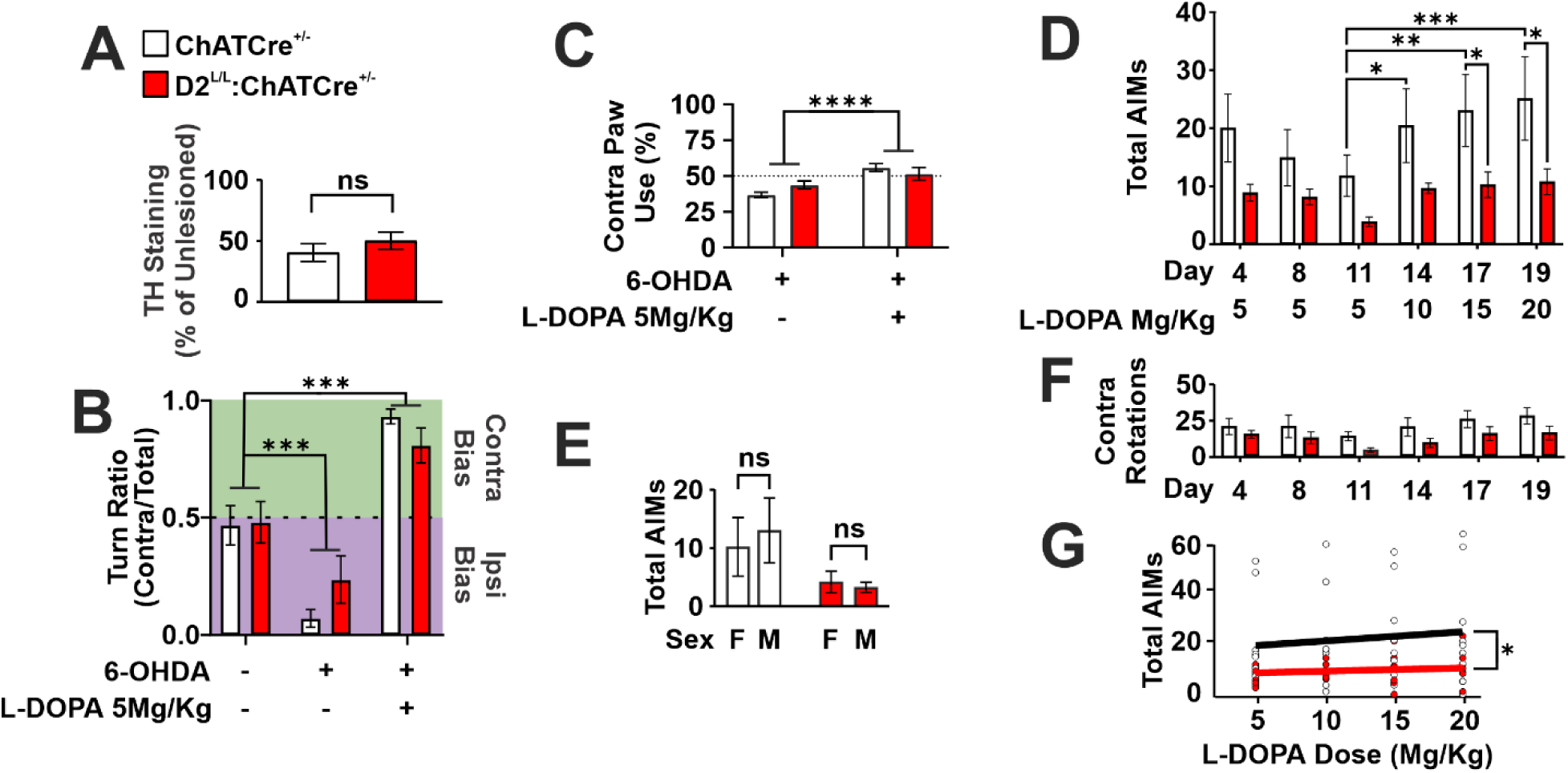
**(A)** Quantification of tyrosine hydroxylase (TH) reduction in the dorsal striatum of animals from the second replication experiment following 6-hydroxydopamine (6-OHDA) injection. Values are expressed as the ratio of lesioned/unlesioned staining intensity in ChATCre^+/-^ controls or D2^L/L^:ChATCre^+/-^ animals (n = 8–9; unpaired two-tailed Student’s t test, p > 0.05). **(B)** Changes in turn bias in animals from the second replication experiment, measured at baseline, after unilateral 6-OHDA lesion, and after subsequent L-DOPA treatment (5Mg/Kg). Turn bias was quantified as the ratio of contralateral turns over total turns (n = 8–9 per genotype; two-way repeated-measures ANOVA: Timepoint effect, F(1.795, 26.92) = 60.24, p < 0.0001; Genotype effect, F(1, 15) = 0.07, p > 0.05; Timepoint × Genotype interaction, F(2, 30) = 2.43, p > 0.05; post hoc Tukey’s multiple comparisons test: ***p < 0.001). **(C)** Contralateral paw use in unilateral 6-OHDA animals from the second replication experiment, measured before and after L-DOPA treatment. Data represent the percentage of total rears performed with the forelimb contralateral to the lesion (n = 8–9 per genotype; two-way repeated-measures ANOVA: Timepoint effect, F(1, 15) = 31.52, p < 0.0001; Genotype effect, F(1, 15) = 0.12, p > 0.05; Timepoint × Genotype interaction, F(1, 15) = 5.59, p < 0.05). **(D)** Replication of total abnormal involuntary movement (AIM) scores across 19 days of L-DOPA treatment (n = 8–9 per genotype; two-way repeated-measures ANOVA: Day/Dose effect, F(5, 75) = 3.98, p < 0.01; Genotype effect, F(1, 15) = 3.92, p > 0.05; Genotype × Day/Dose interaction, F(5, 75) = 0.60, p > 0.05; post hoc Tukey’s multiple comparisons test: *p < 0.05, **p < 0.01, ***p < 0.001). **(E)** Total AIM scores at 5Mg/Kg L-DOPA, grouped by sex and genotype (n = 4–6 per condition; two-way ANOVA: Sex effect, F(1, 13) = 0.05, p > 0.05; Genotype effect, F(1, 13) = 3.62, p > 0.05; Sex × Genotype interaction, F(1, 13) = 0.20, p > 0.05). **(F)** Total contralateral rotations recorded across the same 19 days of L-DOPA treatment shown in Panel (D) (n = 8–9 per genotype; two-way repeated-measures ANOVA: Day/Dose effect, F(2.160, 32.41) = 3.36, p < 0.05; Genotype effect, F(1, 15) = 2.97, p > 0.05; Genotype × Day/Dose interaction, F(5, 75) = 0.14, p > 0.05). **(G)** AIM severity plotted against L-DOPA dose for each genotype in the second replication experiment. Slope differences between genotypes were assessed by ANCOVA (n = 8–9 per genotype; F(1, 4) = 12.13, p < 0.05).

**Supplementary Figure 2.**
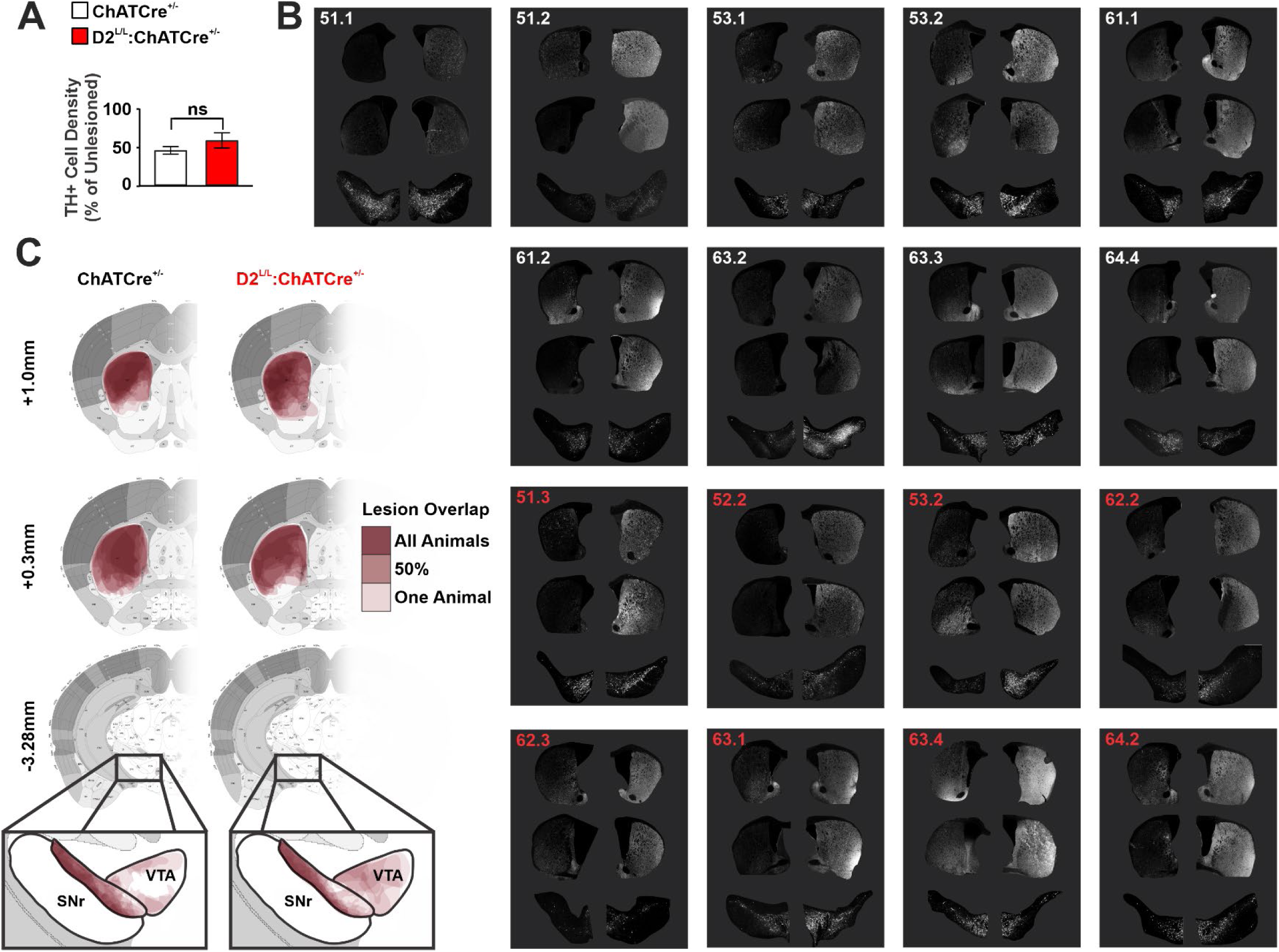
**(A)** Quantification of tyrosine hydroxylase-positive (TH^+^) cell body density in the substantia nigra pars compacta, reported as a percentage of the unlesioned hemisphere (n = 8–9 per genotype; unpaired two-tailed Student’s t test, p > 0.05). **(B)** Representative images of TH immunoreactivity in the striatum and midbrain for all animals included in the replication experiment. For each animal, the left image represents the 6-OHDA-treated hemisphere. ID label color indicates genotype (White = ChATCre^+/-^; Red = D2^L/L^:ChATCre^+/-^). **(C)** Anatomical mapping of 6-OHDA lesion overlap across all animals in each genotype. Regions of TH loss were identified across multiple striatal and midbrain sections by thresholding images to retain the top 10% of pixel intensities. Lesion regions of interest (ROIs) were defined as areas lacking abundant high-intensity TH+ pixels. Individual red contours indicate the ROI of reduced TH intensity for each animal at a given anterior-posterior plane. The color scale reflects the percentage of animals within each cohort showing TH loss at a given anatomical location. ROIs from ChATCre^+/-^ controls are shown on the left, and those from D2^L/L^:ChATCre^+/-^ animals are on the right.

## References

Albin, R. L., A. B. Young, and J. B. Penney. 1989. ‘The functional anatomy of basal ganglia disorders’, Trends Neurosci, 12: 366–75.

Aosaki, T., A. M. Graybiel, and M. Kimura. 1994. ‘Effect of the nigrostriatal dopamine system on acquired neural responses in the striatum of behaving monkeys’, Science, 265: 412–5.

Aosaki, T., H. Tsubokawa, A. Ishida, K. Watanabe, A. M. Graybiel, and M. Kimura. 1994. ’Responses of tonically active neurons in the primate’s striatum undergo systematic changes during behavioral sensorimotor conditioning’, J Neurosci, 14: 3969–84.

Apicella, P., S. Ravel, M. Deffains, and E. Legallet. 2011. ‘The role of striatal tonically active neurons in reward prediction error signaling during instrumental task performance’, J Neurosci, 31: 1507–15.

Bastide, M. F., W. G. Meissner, B. Picconi, S. Fasano, P. O. Fernagut, M. Feyder, V. Francardo, C. Alcacer, Y. Ding, R. Brambilla, G. Fisone, A. Jon Stoessl, M. Bourdenx, M. Engeln, S. Navailles, P. De Deurwaerdere, W. K. Ko, N. Simola, M. Morelli, L. Groc, M. C. Rodriguez, E. V. Gurevich, M. Quik, M. Morari, M. Mellone, F. Gardoni, E. Tronci, D. Guehl, F. Tison, A. R. Crossman, U. J. Kang, K. Steece-Collier, S. Fox, M. Carta, M. Angela Cenci, and E. Bezard. 2015. ’Pathophysiology of L-dopa-induced motor and non-motor complications in Parkinson’s disease’, Progress in neurobiology, 132: 96–168.

Bello, E. P., Y. Mateo, D. M. Gelman, D. Noain, J. H. Shin, M. J. Low, V. A. Alvarez, D. M. Lovinger, and M. Rubinstein. 2011. ‘Cocaine supersensitivity and enhanced motivation for reward in mice lacking dopamine D2 autoreceptors’, Nat Neurosci, 14: 1033–8.

Berke, J. D. 2018. ‘What does dopamine mean?’, Nat Neurosci, 21: 787–93.

Bertran-Gonzalez, J., B. C. Chieng, V. Laurent, E. Valjent, and B. W. Balleine. 2012. ‘Striatal cholinergic interneurons display activity-related phosphorylation of ribosomal protein S6’, PLoS One, 7: e53195.

Bordia, T., X. A. Perez, J. E. Heiss, D. Zhang, and M. Quik. 2016. ‘Optogenetic activation of striatal cholinergic interneurons regulates L-dopa-induced dyskinesias’, Neurobiology of disease, 91: 47–58.

Castello, J., M. Cortes, L. Malave, A. Kottmann, D. R. Sibley, E. Friedman, and H. Rebholz. 2020. ‘The Dopamine D5 receptor contributes to activation of cholinergic interneurons during L-DOPA induced dyskinesia’, Sci Rep, 10: 2542.

Cenci, M. A., H. Jorntell, and P. Petersson. 2018. ‘On the neuronal circuitry mediating L-DOPA-induced dyskinesia’, J Neural Transm (Vienna), 125: 1157–69.

Cenci, M. A., and M. Lundblad. 2007. ’Ratings of L-DOPA-induced dyskinesia in the unilateral 6-OHDA lesion model of Parkinson’s disease in rats and mice’, Curr Protoc Neurosci, Chapter 9: Unit 9 25.

Chantranupong, L., C. C. Beron, J. A. Zimmer, M. J. Wen, W. Wang, and B. L. Sabatini. 2023. ‘Dopamine and glutamate regulate striatal acetylcholine in decision-making’, Nature, 621: 577–85.

Cheung, T. H. C., Y. Ding, X. Zhuang, and U. J. Kang. 2023. ‘Learning critically drives parkinsonian motor deficits through imbalanced striatal pathway recruitment’, Proc Natl Acad Sci U S A, 120: e2213093120.

Choi, S. J., T. C. Ma, Y. Ding, T. Cheung, N. Joshi, D. Sulzer, E. V. Mosharov, and U. J. Kang. 2020. ‘Alterations in the intrinsic properties of striatal cholinergic interneurons after dopamine lesion and chronic L-DOPA’, Elife, 9.

Chuhma, N., S. Mingote, A. Kalmbach, L. Yetnikoff, and S. Rayport. 2017. ‘Heterogeneity in Dopamine Neuron Synaptic Actions Across the Striatum and Its Relevance for Schizophrenia’, Biological psychiatry, 81: 43–51.

Chuhma, N., S. Mingote, H. Moore, and S. Rayport. 2014. ‘Dopamine neurons control striatal cholinergic neurons via regionally heterogeneous dopamine and glutamate signaling’, Neuron, 81: 901–12.

Cotzias, G. C., P. S. Papavasiliou, and R. Gellene. 1969. ‘Modification of Parkinsonism--chronic treatment with L-dopa’, N Engl J Med, 280: 337–45.

Crittenden, J. R., S. Zhai, M. Sauvage, T. Kitsukawa, E. Burguiere, M. Thomsen, H. Zhang, C. Costa, G. Martella, V. Ghiglieri, B. Picconi, K. A. Pescatore, E. M. Unterwald, W. S. Jackson, D. E. Housman, S. B. Caine, D. Sulzer, P. Calabresi, A. C. Smith, D. J. Surmeier, and A. M. Graybiel. 2021. ‘CalDAG-GEFI mediates striatal cholinergic modulation of dendritic excitability, synaptic plasticity and psychomotor behaviors’, Neurobiol Dis, 158: 105473.

Day, M., D. Wokosin, J. L. Plotkin, X. Tian, and D. J. Surmeier. 2008. ‘Differential excitability and modulation of striatal medium spiny neuron dendrites’, The Journal of neuroscience: the official journal of the Society for Neuroscience, 28: 11603–14.

de la Fuente-Fernandez, R., V. Sossi, Z. Huang, S. Furtado, J. Q. Lu, D. B. Calne, T. J. Ruth, and A. J. Stoessl. 2004. ’Levodopa-induced changes in synaptic dopamine levels increase with progression of Parkinson’s disease: implications for dyskinesias’, Brain, 127: 2747–54.

Deffains, M., and H. Bergman. 2015. ‘Striatal cholinergic interneurons and cortico-striatal synaptic plasticity in health and disease’, Mov Disord, 30: 1014–25.

Ding, J. B., J. N. Guzman, J. D. Peterson, J. A. Goldberg, and D. J. Surmeier. 2010. ‘Thalamic gating of corticostriatal signaling by cholinergic interneurons’, Neuron, 67: 294–307.

Ding, J., J. N. Guzman, T. Tkatch, S. Chen, J. A. Goldberg, P. J. Ebert, P. Levitt, C. J. Wilson, H. E. Hamm, and D. J. Surmeier. 2006. ‘RGS4-dependent attenuation of M4 autoreceptor function in striatal cholinergic interneurons following dopamine depletion’, Nature neuroscience, 9: 832–42.

Ding, Y., L. Won, J. P. Britt, S. A. Lim, D. S. McGehee, and U. J. Kang. 2011. ‘Enhanced striatal cholinergic neuronal activity mediates L-DOPA-induced dyskinesia in parkinsonian mice’, Proceedings of the National Academy of Sciences of the United States of America, 108: 840–5.

Duhne, M., A. Mohebi, K. Kim, L. Pelattini, and J. D. Berke. 2024. ‘A mismatch between striatal cholinergic pauses and dopaminergic reward prediction errors’, Proc Natl Acad Sci U S A, 121: e2410828121.

English, D. F., O. Ibanez-Sandoval, E. Stark, F. Tecuapetla, G. Buzsaki, K. Deisseroth, J. M. Tepper, and T. Koos. 2011. ‘GABAergic circuits mediate the reinforcement-related signals of striatal cholinergic interneurons’, Nat Neurosci, 15: 123–30.

Faust, T. W., M. Assous, J. M. Tepper, and T. Koos. 2016. ‘Neostriatal GABAergic Interneurons Mediate Cholinergic Inhibition of Spiny Projection Neurons’, J Neurosci, 36: 9505–11.

Figge, D. A., H. O. Amaral, J. Crim, R. M. Cowell, D. G. Standaert, and K. L. Eskow Jaunarajs. 2024. ‘Differential Activation States of Direct Pathway Striatal Output Neurons during l-DOPA-Induced Dyskinesia Development’, J Neurosci, 44.

Gallo, E. F., J. Greenwald, J. Yeisley, E. Teboul, K. M. Martyniuk, J. M. Villarin, Y. Li, J. A. Javitch, P. D. Balsam, and C. Kellendonk. 2022. ‘Dopamine D2 receptors modulate the cholinergic pause and inhibitory learning’, Mol Psychiatry, 27: 1502–14.

Gerfen, C. R., and D. J. Surmeier. 2011. ‘Modulation of striatal projection systems by dopamine’, Annual review of neuroscience, 34: 441–66.

Girasole, A. E., M. Y. Lum, D. Nathaniel, C. J. Bair-Marshall, C. J. Guenthner, L. Luo, A. C. Kreitzer, and A. B. Nelson. 2018. ‘A Subpopulation of Striatal Neurons Mediates Levodopa-Induced Dyskinesia’, Neuron, 97: 787–95 e6.

Grandy, D. K., Y. A. Zhang, C. Bouvier, Q. Y. Zhou, R. A. Johnson, L. Allen, K. Buck, J. R. Bunzow, J. Salon, and O. Civelli. 1991. ‘Multiple human D5 dopamine receptor genes: a functional receptor and two pseudogenes’, Proc Natl Acad Sci U S A, 88: 9175–9.

Graybiel, A. M., T. Aosaki, A. W. Flaherty, and M. Kimura. 1994. ‘The basal ganglia and adaptive motor control’, Science, 265: 1826–31.

Helseth, A. R., R. Hernandez-Martinez, V. L. Hall, M. L. Oliver, B. D. Turner, Z. F. Caffall, J. E. Rittiner, M. K. Shipman, C. S. King, V. Gradinaru, C. Gerfen, M. Costa-Mattioli, and N. Calakos. 2021. ‘Cholinergic neurons constitutively engage the ISR for dopamine modulation and skill learning in mice’, Science, 372.

Hunker, A. C., M. E. Wirthlin, G. Gill, N. J. Johansen, M. Hooper, V. Omstead, S. Vargas, M. N. Lerma, N. Taskin, N. Weed, W. D. Laird, Y. M. Bishaw, J. L. Bendrick, B. B. Gore, Y. Ben-Simon, X. Opitz-Araya, R. A. Martinez, S. W. Way, B. Thyagarajan, S. Otto, R. E. A. Sanchez, J. R. Alexander, A. Amaya, A. Amster, J. Arbuckle, A. Ayala, P. M. Baker, T. Barcelli, S. Barta, D. Bertagnolli, C. Bielstein, P. Bishwakarma, J. Bowlus, G. Boyer, K. Brouner, B. Casian, T. Casper, A. B. Chakka, R. Chakrabarty, P. Chong, M. Clark, K. Colbert, S. Daniel, T. Dawe, M. Departee, P. DiValentin, N. P. Donadio, N. I. Dotson, D. Dwivedi, T. Egdorf, T. Fliss, A. Gary, J. Goldy, C. Grasso, E. L. Groce, K. Gudsnuk, W. Han, Z. Haradon, S. Hastings, O. Helback, W. V. Ho, C. Huang, T. Johnson, D. L. Jones, Z. Juneau, J. Kenney, M. Leibly, S. Li, E. Liang, H. Loeffler, N. A. Lusk, Z. Madigan, J. Malloy, J. Malone, R. McCue, J. Melchor, J. K. Mich, S. Moosman, E. Morin, R. Naidoo, D. Newman, K. Ngo, K. Nguyen, A. L. Oster, B. Ouellette, A. A. Oyama, N. Pena, T. Pham, E. Phillips, C. Pom, L. Potekhina, S. Ransford, P. L. Ray, M. Reding, D. F. Rette, C. Reynoldson, C. Rimorin, A. R. Sigler, D. B. Rocha, K. Ronellenfitch, A. Ruiz, L. Sawyer, J. P. Sevigny, N. V. Shapovalova, N. Shepard, L. Shulga, S. Soliman, B. Staats, M. J. Taormina, M. Tieu, Y. Wang, J. Wilkes, T. Wood, T. Zhou, A. Williford, N. Dee, T. Mollenkopf, L. Ng, L. Esposito, B. E. Kalmbach, S. Yao, J. Ariza, F. Collman, S. Mufti, K. Smith, J. Waters, I. Ersing, M. Patrick, H. Zeng, E. S. Lein, Y. Kojima, G. Horwitz, S. F. Owen, B. P. Levi, T. L. Daigle, B. Tasic, T. E. Bakken, and J. T. Ting. 2025. ‘Enhancer AAV toolbox for accessing and perturbing striatal cell types and circuits’, Neuron, 113: 1507–24 e17.

Jarvie, K. R., M. Tiberi, C. Silvia, J. A. Gingrich, and M. G. Caron. 1993. ‘Molecular cloning, stable expression and desensitization of the human dopamine D1b/D5 receptor’, J Recept Res, 13: 573–90.

Khan, S. S., E. Jaimon, Y. E. Lin, J. Nikoloff, F. Tonelli, D. R. Alessi, and S. R. Pfeffer. 2024. ’Loss of primary cilia and dopaminergic neuroprotection in pathogenic LRRK2-driven and idiopathic Parkinson’s disease’, Proc Natl Acad Sci U S A, 121: e2402206121.

Khan, S. S., Y. Sobu, H. S. Dhekne, F. Tonelli, K. Berndsen, D. R. Alessi, and S. R. Pfeffer. 2021. ‘Pathogenic LRRK2 control of primary cilia and Hedgehog signaling in neurons and astrocytes of mouse brain’, Elife, 10.

Kharkwal, G., D. Radl, R. Lewis, and E. Borrelli. 2016. ‘Dopamine D2 receptors in striatal output neurons enable the psychomotor effects of cocaine’, Proc Natl Acad Sci U S A, 113: 11609–14.

Kimura, M., J. Rajkowski, and E. Evarts. 1984. ‘Tonically discharging putamen neurons exhibit set-dependent responses’, Proc Natl Acad Sci U S A, 81: 4998–5001.

Koos, T., and J. M. Tepper. 2002. ‘Dual cholinergic control of fast-spiking interneurons in the neostriatum’, J Neurosci, 22: 529–35.

Krok, A. C., M. Maltese, P. Mistry, X. Miao, Y. Li, and N. X. Tritsch. 2023. ‘Intrinsic dopamine and acetylcholine dynamics in the striatum of mice’, Nature, 621: 543–49.

Laverne, G., J. Pesce, A. Reynders, E. Combrisson, E. Gascon, C. Melon, L. Kerkerian-Le Goff, N. Maurice, and C. Beurrier. 2022. ‘Cholinergic interneuron inhibition potentiates corticostriatal transmission in direct medium spiny neurons and rescues motor learning in parkinsonism’, Cell Rep, 40: 111034.

Lim, S. A., U. J. Kang, and D. S. McGehee. 2014. ‘Striatal cholinergic interneuron regulation and circuit effects’, Frontiers in synaptic neuroscience, 6: 22.

Longo, F., S. Aryal, P. G. Anastasiades, M. Maltese, C. Baimel, F. Albanese, J. Tabor, J. D. Zhu, M. M. Oliveira, D. Gastaldo, C. Bagni, E. Santini, N. X. Tritsch, A. G. Carter, and E. Klann. 2023. ‘Cell-type-specific disruption of cortico-striatal circuitry drives repetitive patterns of behavior in fragile X syndrome model mice’, Cell Rep, 42: 112901.

Lucarelli, M., C. Di Pietro, G. La Sala, M. T. Fiorenza, D. Marazziti, and S. Canterini. 2019. ‘Anomalies in Dopamine Transporter Expression and Primary Cilium Distribution in the Dorsal Striatum of a Mouse Model of Niemann-Pick C1 Disease’, Front Cell Neurosci, 13: 226.

Malave, L., D. R. Zuelke, S. Uribe-Cano, L. Starikov, H. Rebholz, E. Friedman, C. Qin, Q. Li, E. Bezard, and A. H. Kottmann. 2021. ’Dopaminergic co-transmission with sonic hedgehog inhibits abnormal involuntary movements in models of Parkinson’s disease and L-Dopa induced dyskinesia’, Commun Biol, 4: 1071.

Marley, A., and M. von Zastrow. 2010. ‘DISC1 regulates primary cilia that display specific dopamine receptors’, PLoS One, 5: e10902.

Martyniuk, K. M., A. Torres-Herraez, D. C. Lowes, M. Rubinstein, M. A. Labouesse, and C. Kellendonk. 2022. ‘Dopamine D2Rs coordinate cue-evoked changes in striatal acetylcholine levels’, Elife, 11.

Matamales, M., J. Gotz, and J. Bertran-Gonzalez. 2016. ‘Quantitative Imaging of Cholinergic Interneurons Reveals a Distinctive Spatial Organization and a Functional Gradient across the Mouse Striatum’, PLoS One, 11: e0157682.

Miyoshi, K., K. Kasahara, S. Murakami, M. Takeshima, N. Kumamoto, A. Sato, I. Miyazaki, S. Matsuzaki, T. Sasaoka, T. Katayama, and M. Asanuma. 2014. ‘Lack of dopaminergic inputs elongates the primary cilia of striatal neurons’, PLoS One, 9: e97918.

Morris, G., D. Arkadir, A. Nevet, E. Vaadia, and H. Bergman. 2004. ‘Coincident but distinct messages of midbrain dopamine and striatal tonically active neurons’, Neuron, 43: 133–43.

Nair, S. V., E. Jaimon, A. Adhikari, J. Nikoloff, and S. R. Pfeffer. 2025. ’Lysosomal glucocerebrosidase is needed for ciliary Hedgehog signaling: A convergent pathway contributing to Parkinson’s disease’, Proc Natl Acad Sci U S A, 122: e2504774122.

Nielsen, B. E., and C. P. Ford. 2024. ‘Reduced striatal M4-cholinergic signaling following dopamine loss contributes to parkinsonian and l-DOPA-induced dyskinetic behaviors’, Sci Adv, 10: eadp6301.

Oldenburg, I. A., and J. B. Ding. 2011. ‘Cholinergic modulation of synaptic integration and dendritic excitability in the striatum’, Curr Opin Neurobiol, 21: 425–32.

Pisani, A., G. Martella, A. Tscherter, P. Bonsi, N. Sharma, G. Bernardi, and D. G. Standaert. 2006. ‘Altered responses to dopaminergic D2 receptor activation and N-type calcium currents in striatal cholinergic interneurons in a mouse model of DYT1 dystonia’, Neurobiol Dis, 24: 318–25.

Ratna, D. D., and T. C. Francis. 2025. ‘Extrinsic and intrinsic control of striatal cholinergic interneuron activity’, Front Mol Neurosci, 18: 1528419.

Ravel, S., E. Legallet, and P. Apicella. 1999. ‘Tonically active neurons in the monkey striatum do not preferentially respond to appetitive stimuli’, Exp Brain Res, 128: 531–4.

Redgrave, P., M. Rodriguez, Y. Smith, M. C. Rodriguez-Oroz, S. Lehericy, H. Bergman, Y. Agid, M. R. DeLong, and J. A. Obeso. 2010. ’Goal-directed and habitual control in the basal ganglia: implications for Parkinson’s disease’, Nature reviews. Neuroscience, 11: 760–72.

Rossi, J., N. Balthasar, D. Olson, M. Scott, E. Berglund, C. E. Lee, M. J. Choi, D. Lauzon, B. B. Lowell, and J. K. Elmquist. 2011. ‘Melanocortin-4 receptors expressed by cholinergic neurons regulate energy balance and glucose homeostasis’, Cell Metab, 13: 195–204.

Ryan, M. B., A. E. Girasole, A. J. Flores, E. L. Twedell, M. M. McGregor, R. Brakaj, R. F. Paletzki, T. S. Hnasko, C. R. Gerfen, and A. B. Nelson. 2024. ‘Excessive firing of dyskinesia-associated striatal direct pathway neurons is gated by dopamine and excitatory synaptic input’, Cell Rep, 43: 114483.

Schulz, J. M., and J. N. Reynolds. 2013. ‘Pause and rebound: sensory control of cholinergic signaling in the striatum’, Trends in neurosciences, 36: 41–50.

Sebastianutto, I., N. Maslava, C. R. Hopkins, and M. A. Cenci. 2016a. ‘Validation of an improved scale for rating l-DOPA-induced dyskinesia in the mouse and effects of specific dopamine receptor antagonists’, Neurobiology of disease, 96: 156–70.

Sebastianutto, I., N. Maslava, C. R. Hopkins, and M. A. Cenci. 2016b. ‘Validation of an improved scale for rating l-DOPA-induced dyskinesia in the mouse and effects of specific dopamine receptor antagonists’, Neurobiol Dis, 96: 156–70.

Shen, W., J. L. Plotkin, V. Francardo, W. K. Ko, Z. Xie, Q. Li, T. Fieblinger, J. Wess, R. R. Neubig, C. W. Lindsley, P. J. Conn, P. Greengard, E. Bezard, M. A. Cenci, and D. J. Surmeier. 2016. ‘M4 Muscarinic Receptor Signaling Ameliorates Striatal Plasticity Deficits in Models of L-DOPA-Induced Dyskinesia’, Neuron, 90: 1139.

Shen, W., X. Tian, M. Day, S. Ulrich, T. Tkatch, N. M. Nathanson, and D. J. Surmeier. 2007. ‘Cholinergic modulation of Kir2 channels selectively elevates dendritic excitability in striatopallidal neurons’, Nature neuroscience, 10: 1458–66.

Shen, W., S. Zhai, and D. J. Surmeier. 2022. ’Striatal synaptic adaptations in Parkinson’s disease’, Neurobiol Dis, 167: 105686.

Straub, C., N. X. Tritsch, N. A. Hagan, C. Gu, and B. L. Sabatini. 2014. ‘Multiphasic modulation of cholinergic interneurons by nigrostriatal afferents’, The Journal of neuroscience: the official journal of the Society for Neuroscience, 34: 8557–69.

Sullivan, M. A., H. Chen, and H. Morikawa. 2008. ‘Recurrent inhibitory network among striatal cholinergic interneurons’, J Neurosci, 28: 8682–90.

Surmeier, D. J., L. Carrillo-Reid, and J. Bargas. 2011. ‘Dopaminergic modulation of striatal neurons, circuits, and assemblies’, Neuroscience, 198: 3–18.

Surmeier, D. J., J. Ding, M. Day, Z. Wang, and W. Shen. 2007. ‘D1 and D2 dopamine-receptor modulation of striatal glutamatergic signaling in striatal medium spiny neurons’, Trends in neurosciences, 30: 228–35.

Tiberi, M., and M. G. Caron. 1994. ‘High agonist-independent activity is a distinguishing feature of the dopamine D1B receptor subtype’, J Biol Chem, 269: 27925–31.

Tiberi, M., K. R. Jarvie, C. Silvia, P. Falardeau, J. A. Gingrich, N. Godinot, L. Bertrand, T. L. Yang-Feng, R. T. Fremeau, Jr., and M. G. Caron. 1991. ‘Cloning, molecular characterization, and chromosomal assignment of a gene encoding a second D1 dopamine receptor subtype: differential expression pattern in rat brain compared with the D1A receptor’, Proc Natl Acad Sci U S A, 88: 7491–5.

Tiberi, M., S. R. Nash, L. Bertrand, R. J. Lefkowitz, and M. G. Caron. 1996. ‘Differential regulation of dopamine D1A receptor responsiveness by various G protein-coupled receptor kinases’, J Biol Chem, 271: 3771–8.

Tritsch, N. X., J. B. Ding, and B. L. Sabatini. 2012. ‘Dopaminergic neurons inhibit striatal output through non-canonical release of GABA’, Nature, 490: 262–6.

Tritsch, N. X., and B. L. Sabatini. 2012. ‘Dopaminergic modulation of synaptic transmission in cortex and striatum’, Neuron, 76: 33–50.

Tubert, C., R. M. Paz, A. M. Stahl, K. M. Sanchez Armijos, L. Rela, and M. G. Murer. 2025. ‘Striatal cholinergic interneuron pause response requires Kv1 channels, is absent in dyskinetic mice, and is restored by dopamine D5 receptor inverse agonism’, Elife, 13.

Uribe-Cano, S., and A. H. Kottmann. 2024. ’The primary cilium of cholinergic neurons may be a linchpin in the progression of Parkinson’s Disease’, Proc Natl Acad Sci U S A, 121: e2414226121.

Uribe-Cano, S., and A. H. Kottmann. 2025. ‘The GPCR Smoothened on Cholinergic Interneurons Modulates Dopamine-associated Acetylcholine Dynamics’, bioRxiv: 2025.07.03.662982.

Wang, Z., L. Kai, M. Day, J. Ronesi, H. H. Yin, J. Ding, T. Tkatch, D. M. Lovinger, and D. J. Surmeier. 2006. ‘Dopaminergic control of corticostriatal long-term synaptic depression in medium spiny neurons is mediated by cholinergic interneurons’, Neuron, 50: 443–52.

Wieland, S., D. Du, M. J. Oswald, R. Parlato, G. Kohr, and W. Kelsch. 2014. ‘Phasic dopaminergic activity exerts fast control of cholinergic interneuron firing via sequential NMDA, D2, and D1 receptor activation’, J Neurosci, 34: 11549–59.

Wieland, S., S. Schindler, C. Huber, G. Kohr, M. J. Oswald, and W. Kelsch. 2015. ‘Phasic Dopamine Modifies Sensory-Driven Output of Striatal Neurons through Synaptic Plasticity’, The Journal of neuroscience: the official journal of the Society for Neuroscience, 35: 9946–56.

Won, L., Y. Ding, P. Singh, and U. J. Kang. 2014. ‘Striatal cholinergic cell ablation attenuates L-DOPA induced dyskinesia in Parkinsonian mice’, The Journal of neuroscience: the official journal of the Society for Neuroscience, 34: 3090–4.

Yan, Z., W. J. Song, and J. Surmeier. 1997. ‘D2 dopamine receptors reduce N-type Ca2+ currents in rat neostriatal cholinergic interneurons through a membrane-delimited, protein-kinase-C-insensitive pathway’, J Neurophysiol, 77: 1003–15.

Yan, Z., and D. J. Surmeier. 1997. ‘D5 dopamine receptors enhance Zn2+-sensitive GABA(A) currents in striatal cholinergic interneurons through a PKA/PP1 cascade’, Neuron, 19: 1115–26.

Zhai, H., Z. Kang, H. Zhang, J. Ma, and G. Chen. 2019. ’Baicalin attenuated substantia nigra neuronal apoptosis in Parkinson’s disease rats via the mTOR/AKT/GSK-3beta pathway’, J Integr Neurosci, 18: 423–29.

Zhai, S., Q. Cui, D. V. Simmons, and D. J. Surmeier. 2023. ’Distributed dopaminergic signaling in the basal ganglia and its relationship to motor disability in Parkinson’s disease’, Curr Opin Neurobiol, 83: 102798.

Zhai, S., Q. Cui, D. Wokosin, L. Sun, T. Tkatch, J. R. Crittenden, A. M. Graybiel, and D. J. Surmeier. 2025. ’State-dependent modulation of spiny projection neurons controls levodopa-induced dyskinesia in a mouse model of Parkinson’s disease’, Sci Adv, 11: eadv8224.

Zhai, S., A. Tanimura, S. M. Graves, W. Shen, and D. J. Surmeier. 2017. ’Striatal synapses, circuits, and Parkinson’s disease’, Current opinion in neurobiology, 48: 9–16.

Zhang, Y. F., and S. J. Cragg. 2017. ‘Pauses in Striatal Cholinergic Interneurons: What is Revealed by Their Common Themes and Variations?’, Front Syst Neurosci, 11: 80.

Zhuang, X., P. Mazzoni, and U. J. Kang. 2013. ‘The role of neuroplasticity in dopaminergic therapy for Parkinson disease’, Nature reviews. Neurology, 9: 248–56.

